# Mass spectrometry-based quantitation of Her2 in gastroesophageal tumor tissue: Comparison to IHC and FISH

**DOI:** 10.1101/018747

**Authors:** Daniel V.T. Catenacci, Wei-Li Liao, Lei Zhao, Emma Whitcomb, Les Henderson, Emily O’Day, Peng Xu, Sheeno Thyparambil, David Krisman, Kathleen Bengali, Jamar Uzzell, Marlene Darflur, Fabiola Cecchi, Adele Blackler, Yung-Jue Bang, John Hart, Shu-Yuan Xiao, Sang Mee Lee, Jon Burrows, Todd Hembrough

**Affiliations:** University of Chicago, Department of Medicine, Section of Hematology & Oncology, Chicago, IL; OncoPlex Diagnostics Inc., Rockville, MD; University of Chicago, Department of Pathology, Chicago, IL; Seoul National University College of Medicine, Seoul, Korea; University of Chicago, Department of Health Studies, Chicago, IL

**Keywords:** Her2 expression, *HER2* (*ERBB2*) amplification, gastric, esophageal, gastroesophageal adenocarcinoma, stomach cancer, SRM-MS, selected reaction monitoring mass spectrometry, companion diagnostic, clinical biomarker assay, multiplex protein expression analysis in FFPE tissue

## Abstract

**Background:** Trastuzumab showed survival benefit for Her2-positive gastroesophageal cancers (GEC). Immunohistochemistry (IHC) and fluorescence *in situ* hybridization (FISH) currently determine eligibility for trastuzumab-based therapy. However, both assays are low throughput with various limitations.

**Methods:** We developed a selected reaction monitoring mass spectrometric (SRM-MS) assay and quantified levels (amol/ug) of Her2-SRM in cell lines (n=27) and GEC tissues (n=139). We compared Her2-SRM expression with IHC/FISH, seeking to determine optimal SRM expression cut-offs to identify *HER2* amplification.

**Results:** After demonstrating assay development, precision, and stability, Her2-SRM measurement was observed to be highly concordant with *HER2/CEP17* ratio, particularly in a multivariate regression model adjusted for SRM-expression of Met, Egfr, Her3, and HER2-heterogeneity covariates, and their interactions (cell lines r^2^=0.9842; FFPE r^2^=0.7643). In GEC tissues, Her2-SRM was detected in 71.2% of cases, and 12.3% were identified as ‘HER2+’. ROC curves demonstrated HER2-SRM levels to have high specificity (100%) at an upper-level cut-off of **>**750 amol/μg and sensitivity (75%) at lower-level cut-off of <450 amol/ug. We observed an ‘equivocal-zone’ between 450-750 amol/ug, analogous to ‘IHC2+’, but less frequent (9-16% of cases versus 36-41%).

**Significance:** Compared to IHC, SRM-MS provided more objective and quantitative Her2 expression with excellent *HER2/CEP17* FISH correlation and fewer ‘equivocal’ cases. Along with the multiplex capability for other relevant oncoproteins, these results demonstrated a refined HER2 expression assay for clinical application.

## Introduction

The human epidermal growth factor receptor-2 (HER2, *ERBB2*) is a receptor tyrosine kinase promoting cell development, differentiation and survival.^1, 2^ Aberrant HER2 activity due to gene amplification and consequent protein overexpression results in a HER2-driven oncogenic phenotype.^1, 2^ HER2 is amplified/overexpressed in various cancers including breast (∼20%), gastroesophageal (GEC) (∼10-15%) and endometrial cancers (∼12%).^3^ HER2 positivity is higher in esophageal/esophagogastric adenocarcinomas (∼15%), compared to distal gastric adenocarcinomas (∼10%).^4, 5^ The ‘**T**rastuzumab in the treatment **O**f **GA**stric cancer’ (‘ToGA’) trial reported survival benefit among ‘HER2-positive’ GEC patients treated with trastuzumab-based therapy in comparison to standard chemotherapy, with ensuing widespread incorporation of immunohistochemistry (IHC) and/or fluorescence in situ hybridization (FISH) testing into routine GEC care (**Supplementary Figure 1**).^4^

‘ToGA’ trial eligibility defined ‘HER2 positivity’ as either ‘FISH+/any IHC’ or ‘anyFISH/IHC3+’. However, patients within ‘FISH+/IHC 0-1+’ subgroups (n=131 or 22% of enrolled patients) derived no benefit from trastuzumab (**Supplementary Figure 2**). In ‘ToGA’, FISH positivity was defined as ‘*HER2/CEP17* ratio of ≥2’. IHC validation on GEC samples led to modification of the breast scoring system to account for basolateral membranous immunoreactivity and/or higher rates of intra-tumoral heterogeneity[6]. Now, ‘HER2+’ is clinically defined as IHC3+ or IHC2+/FISH+ **(Supplementary Figure 1,2)**.^6^-^9^ Subsequent phase III “HER2-selective” GEC trials in the first line (‘LOGiC’)^10^ and second line (‘TyTAN’)^11^ metastatic settings evaluated the HER2/Egfr specific oral tyrosine kinase inhibitor, lapatinib, versus placebo along with chemotherapy; both trials were negative for the primary endpoint (overall survival) in the intention-to-treat populations. Interestingly, ‘TyTAN’ enrolled 261 patients of which 31% were FISH+/IHC 0-1+.^11^ Moreover, the IHC3+/FISH+ subset demonstrated a statistically significant survival advantage (14 vs 7.6 months, HR 0.59, p=0.0175).^11^ Recently, a report suggested that the degree of *HER2* amplification/expression may better predict therapeutic benefit from anti-HER2 therapy.^12^ These observations suggest the need for revised HER2 criteria/diagnostics, and also implications regarding optimal therapeutic strategies *within* classic ‘HER2+’ groups.^13, 14^

Despite noted utility of HER2 IHC/FISH, various reports detail numerous limitations.^15^-^20^ IHC is semi-quantitative attempting to incorporate staining intensity and extensity into a ‘0-3+’ scoring system. IHC is notoriously subjective, and sensitive to antigen instability in formalin fixed paraffin embedded (FFPE) unstained-sections, as recently demonstrated.^21-24^ ‘HER2-equivocal’ (IHC2+) scores require reflex FISH analysis – accounting for almost 30% (159/584) of FISH+ enrolled cases in ‘ToGA’ (**Supplementary Figure 2A**), not including undocumented IHC2+/FISH-screen failures. Reflex FISH testing is also laborious, time-consuming (especially serially after IHC), expensive, and remains operator dependent/subjective, particularly in molecularly heterogeneous cases.^7, 25-28^ Both assays are low throughput.^13^ A consequence of these limitations is false positive/ negative and delayed results and uneconomical use of limited tissue samples.^15, 19, 26^ Refinement of HER2 diagnostic methods is welcomed.

The mass spectrometry-based selected-reaction-monitoring (SRM-MS) assay has gained broad acceptance as a specific and sensitive technology for measuring absolute levels of specific protein targets,^29-31^ however the ability to apply this technology to FFPE tissues has been technically challenging. A multiplexed and quantitative Liquid Tissue-SRM method to quantify proteins in FFPE tissues based on unique peptide sequences does not have the same technical limitations as IHC/FISH, as recently described.^21-24^

We sought to evaluate a clinical role of Her2-SRM expression for GEC. Herein we describe application of the Her2-SRM assay to 27 cell lines and uniquely to 139 FFPE GEC tumors. We first tested precision and temporal reproducibility of the assay. We next assessed correlation of Her2-SRM with Her2 IHC/FISH scores, along with multivariate modeling accounting for HER2-heterogeneity and SRM-expression of other relevant GEC oncoproteins Met, Egfr, and Her3. We determined optimal Her2-SRM expression cut-off values for clinical application using ROC curves correlating with *HER2/CEP17* FISH ratio. Finally, we present clinical cases to substantiate advantages of the ‘GEC-plex’ assay with the capability of quantifying multiple oncoproteins simultaneously, addressing issues surrounding the current molecular profiling hurdles of inter- and intra-patient molecular heterogeneity.

## Materials and Methods

### GEC Clinical Samples and Cell Lines

Patient samples and cell lines were obtained and processed from the University of Chicago (Chicago, IL) under preapproved protocols as previously described.^23, 32^

Cell line mixing studies with *HER2* amplified OE-19 and *HER2* non-amplified MKN-1 to demonstrate dilutional effects of molecular subclones were performed in six lysate conditions with varying ratios (0/100, 20/80, 40/60, 60/40, 80/20, 100/0).

### Sample Preparation and Her2-SRM Assay Development

Laser microdissection isolated cells were obtained from FFPE tumor sections as previously described.^21-24^ Total protein content for lysates was measured using Micro-BCA assay (Thermo Fisher Scientific Inc, Rockford, IL). Her2-SRM assay development followed previously described methods.^21-24^ Briefly, recombinant Her2 protein (UniProtKB accession number P04626) was digested with trypsin and the resultant peptides analyzed using a TSQ Vantage triple quadrupole mass spectrometer (Thermo Scientific, San Jose, CA) equipped with a nanoAcquityLC system (Waters, Milford, MA). Peptides containing methionine or cysteine-residues were excluded due to their propensity to undergo unpredictable oxidation. Tryptic peptides analysis based on reproducible peak heights, retention times, chromatographic ion intensities, and distinctive/reproducible transition ion ratios identified the optimal SRM peptide ELVSEFSR, comprising residues 971-978 within the protein’s intracellular domain, to be unique to Her2. Light (ELVSEFSR) and heavy (ELVSEFSR [^13^C_6_,^15^N_4_]) versions of this peptide were synthesized to develop and perform the assay (Thermo Scientific, San Jose, CA). SRM transitions, chromatography and mass spectrometer conditions used for the quantification were specified in our previous report^21^

### Her2-SRM Assay Precision & Temporal Reproducibility

Assay precision and temporal stability of the Her2-SRM assay was performed as previously described.^23, 24^ To demonstrate assay precision, eight breast cancer and eleven GEC FFPE tissues were analyzed in triplicate independently by two different operators on two platforms (“System R” and “System S”) on different days, blinded to the others’ results. “System R” was comprised of a nanoAcquity LC coupled to a TSQ Vantage mass spectrometer and “System S” was comprised of a Thermo Easy nLC II coupled to a separate TSQ Vantage mass spectrometer.

Stability of the MS-based SRM-HER2 assay in FFPE tumor tissue was assessed. Serial tissue sections were cut from eighteen GEC tumor blocks and 11 NSCLC tumor blocks. LM and Liquid Tissue preparation were immediately performed on freshly cut FFPE sections then SRM-HER2 performed. Approximately 13 months later processing of the serial tissue sections was performed and SRM-HER2 levels compared to the previous measurements.

### Quantitative Analysis and Validation of Her2 in Clinical GEC Tissues and Cell lines

Her2-SRM for 139 GEC FFPE samples and 27 cell lines was calculated from the ratio of area under the curve (AUC) for the endogenous and isotopically-labeled standard peptide multiplied by the known amount of isotopically-labeled standard peptide spiked into the sample before analysis, as previously described.^21-24^

### HER2 Fluorescence in situ hybridization (FISH)

FISH results were obtained through routine clinical testing for *HER2/CEP17* ratio. The majority of samples which had clinical FISH testing were those with an initial Her2 IHC 2+ score, per routine standards. FISH was retrospectively performed on available IHC0/1/3+ samples missing FISH results, as previously described.^32, 33^

#### *HER2* Heterogeneity

FISH *HER2* heterogeneity (hetero+) was defined as 10-50% of enumerated nuclei having *HER2/CEP17* ratio ≥2.^26^ HER2 negative (HER2-) was defined as <10% of scored nuclei having ratio ≥2, and HER2+ (non-heterogeneous) was defined as >50% of scored nuclei having ratio ≥2.^26^

### Her2 Immunohistochemistry (IHC)

IHC Her2 scores were obtained per routine clinical care using the Hercept Test kit from DAKO.^6^ (**Supplementary Figure 1,2**). For samples without clinical Her2 IHC (eg. archived curative-intent resections), when tissue was available, DAB-labeled dextrose-based polymer complex bound to secondary antibody (Leica Microsystems Inc., Buffalo Grove, IL) was performed as previously described.^34^

### Statistical Methods

To examine association between Her2-SRM and *HER2* gene copy number (GCN) or *HER2/CEP17* ratio in cell lines and tissues, we used univariate and multivariate linear regression models with Her2-SRM as the independent and *HER2* GCN or *HER2/CEP17* ratio as the dependent variable. In multivariate models, we included SRM expression for Met, Egfr, and Her3, and their interaction terms, given their putative association with HER2 signaling. Analyzing the tissue sample data, *HER2* FISH heterogeneity was additionally included in the models. To compare IHC to either FISH or Her2-SRM in GEC tissues, we included indicators for IHC2+ and IHC3+ (IHC0/1+ reference category) in the regression model. To assess the most effective cutoff value for Her2-SRM to identify *HER2/CEP17* ratio, we computed a receiver-operating-characteristic (ROC) curve. All analyses were performed using R software (www.r-project.ort), version 3.0.1.

## Results

### SRM Assay Development

For development of the Her2 Liquid Tissue-SRM assay, multiple peptides obtained from a tryptic digest of recombinant Her2 were measured using MS. The resulting three candidate peptides, SLTEILK, VLQGLPR, and ELVSEFSR, were then extensively screened in multiple formalin-fixed cell lines and FFPE clinical samples. The peptide ELVSEFSR provided the most reproducible peak heights, retention times, chromatographic ion intensities, clean elution profile, and distinctive/reproducible transition ion ratios; therefore, this peptide was selected for clinical assay development. SRM transitions used for the quantification of Her2 was selected based on the representative MS fragmentation spectrum for peptide ELVSEFSR [^13^C_6_,^15^N_4_] (**Figure 1A**). A calibration curve was built in formalin-fixed PC3 cell lysates to assess the linearity and the limits of detection and quantitation (LOD, LOQ) of the assay (**Figure 1B**). Coefficients of variation (CV’s) ranged from 2.20-14.04% for samples (5 replicates). The LOD/LOQ was 150/200 amol, respectively, with a linear regression value of r^2^ = 0.9998. The linearity and tight %CV over the concentration range demonstrated the accuracy, reproducibility, and quantitative resolution of the assay. The total ion chromatograms for the light/heavy isotopically-labeled peptides (**Figure 1C)**, and the transition ions (**Figure 1D**) are shown. SRM-Fgfr2 and SRM-Her3 are detailed in **Supplementary Figure 3**.

**Figure 1.**
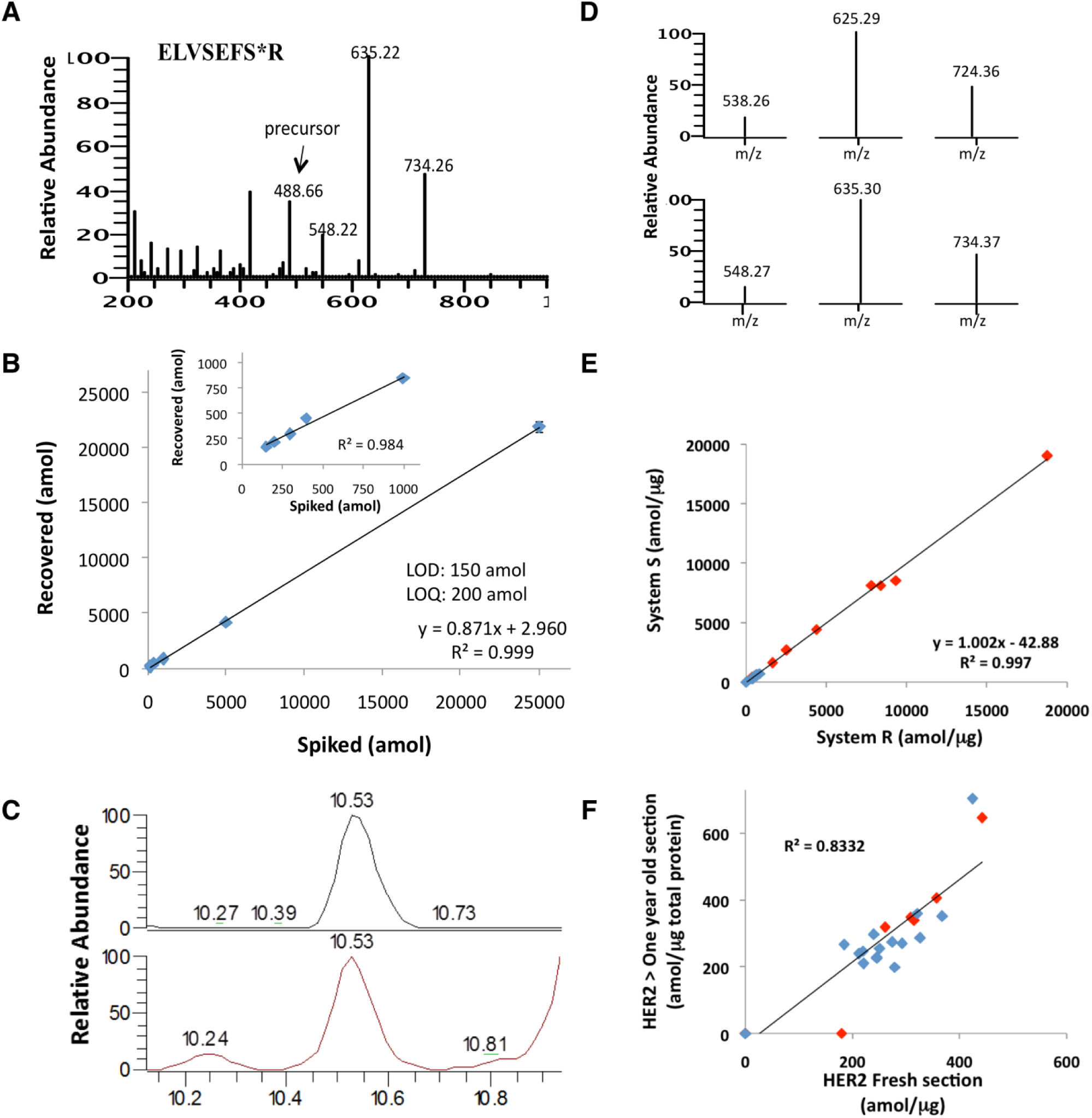
Development of HER2 SRM assay. **(A)** The fragmentation spectrum for heavy ELVSEFSR peptide and **(B)** the standard curve generated in human PC3 cell lysate; inset: the standard curve generated without the highest two spiking points (5000 and 25000 amol). Each point injected contained 5000 amol heavy (ELVSEFSR [^13^C_6_,^15^N_4_]) on column. **(C)** The total ion chromatograms for the light and heavy isotopically labeled peptides, with **(D)** the transition ions used to identify and quantitate each peptide. **(E)** Precision assessment for measuring Her2 level in 8 breast cancer (red) and 11 GEC (blue) FFPE tissues. (**F)** Temporal reproducibility of FFPE sections processed and analyzed using LT-SRM at two time points over one year apart (blue, GEC (n=18); red, NSCLC (n=9)).

### Precision of HER2-SRM Assay

To test assay precision, Her2 was measured in eight human breast cancer and eleven human GEC tissues. Using two different LC-MS systems and operators, all breast cancer samples expressed Her2, ranging 395.1-18896.7 amol/ug, CVs ranging 3.7-10.4%. Nine of eleven GEC samples expressed Her2 (≥LOD) ranging 306-767.7 amol/ug, CVs ranging 7.5-14.6%. The two operating systems showed very good concordance, r^2^ = 0.9978, as demonstrated previously (**Figure 1E, Supplemental Table 1**).**^21, 23^**

### Temporal Reproducibility of Her2-SRM Assay

To test the assay’s temporal reproducibility, two sections from 18 GEC and 9 NSCLC samples were processed 13 months apart. Very good correlation (r^2^=0.8332) supported the reproducibility of Her2-SRM results in archival FFPE sections **(Figure 1F, Supplemental Table 2).**

### Her2-SRM Correlation with FISH in Cell Lines & HER2-heterogeneity Mixing Studies

The correlation of Her2-SRM expression and *HER2* GCN or ratio was assessed in 27 GEC and breast cancer lines. (**Figure 2, Supplemental Table 3).** Her2-SRM ranged from <150-21896.7 amol/ug (**Figure 2A**). Her2**-**SRM results correlated well with *HER2* GCN and ratio in univariate analyses (r^2^ = 0.6096 and 0.7493, respectively) (**Figure 2B,C**). Adjusting for SRM-expressions for Met, Egfr, and Her3, the multivariate regression model resulted in an improved and very good correlation between Her2-SRM and *HER2* GCN or ratio (r^2^ = 0.8829 and 0.9842, respectively) (Supplementary **Figure 4**, **Supplementary Table 5**). Using a preliminary cut-off of 1175 amol/ug, derived from these data, Her2-SRM discerned cell lines with *HER2* amplification versus non-amplification with 100% sensitivity (5/5) and specificity (22/22).

**Figure 2.**
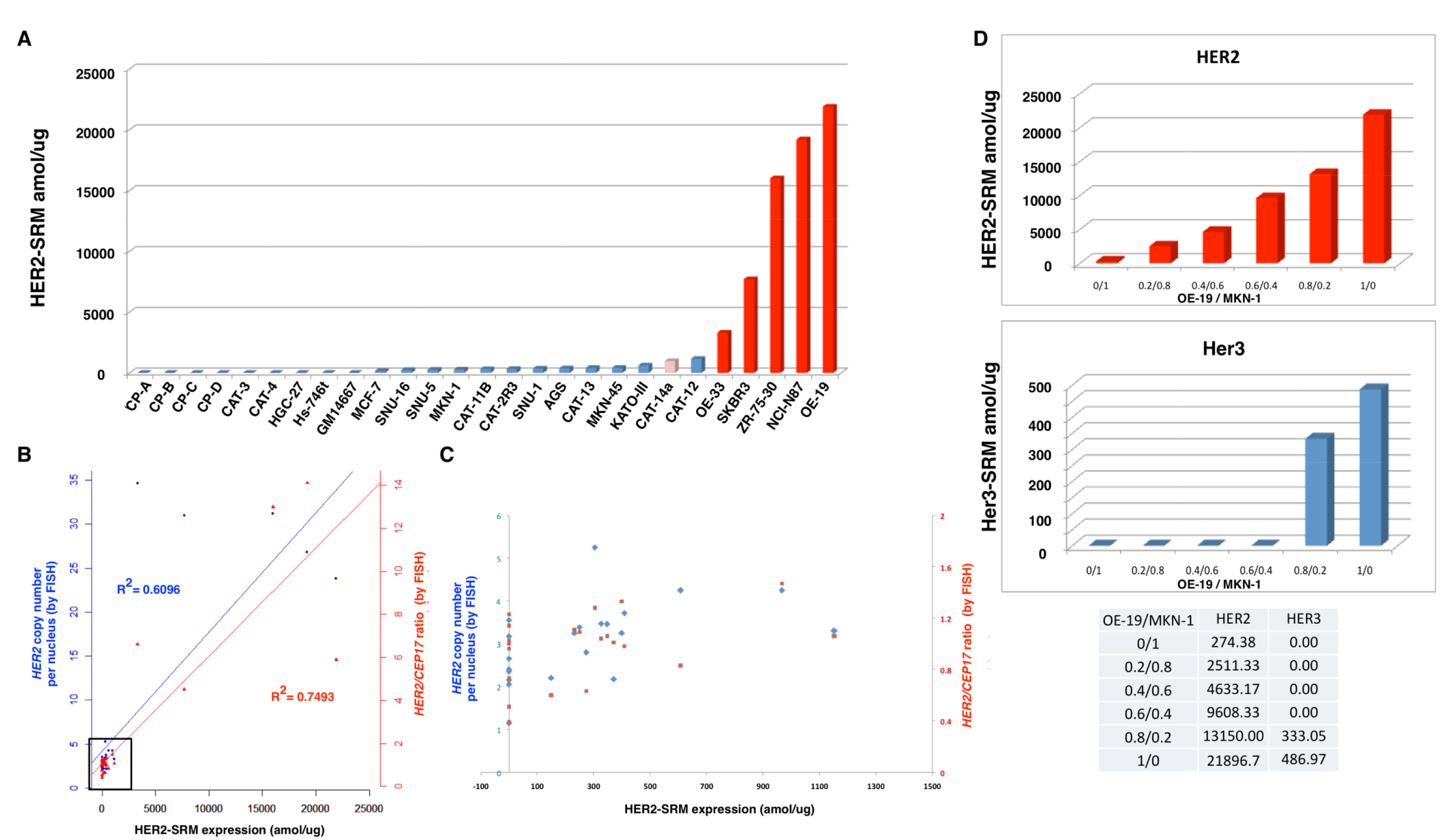
Her2 expression levels using LT-SRM and correlation with *HER2* gene amplification (by FISH) in 27 cell lines. **(A)** Quantification of Her2-SRM (amol/ug) for 27 cell lines including GM15677 lymphoblast control. *HER2* amplified cell lines (*HER2/CEP7* ratio ≥2) indicated in dark red, and heterogenous *HER2* amplification in pink (see methods for hetero+ scoring). **(B)** Her2-SRM and FISH gene copy number (GCN) univariate correlations: The left y-axis (blue plus sign) represents the mean *HER2* GCN per nucleus and the right y-axis (red triangle) indicates *HER2:CEP17* ratio (see text for multivariate analysis including Egfr-, Her3-, and Met-SRM coexpression, GCN R^2^=0.8829 and Ratio R^2^= 0.9824) and **(C)** scatter plot of *HER2* GCN/ratio (by FISH) (blue diamond, red square, respectively) and Her2-SRM expression in samples where Her2 expressions are < 1500 amol/ug represented by the black box in (3B). A preliminary Her2-SRM cut-off, from these cell line data, correlating with *HER2* amplification was determined to be ≥1150 amol/ug. **(D)** Her2-SRM (red) and Her3-SRM (blue) levels in a cell line mixing study (OE-19: *HER2* amplified / MKN-1: *HER2* non-amplified), modeling intra-tumor clonal heterogeneity.

One cell line, CAT-14a, was observed to be ‘HER2+/hetero+’, having 30% of nuclei with ratio ≥2; Her2-SRM level was 969.33amol/ug, below the established cut-off of 1175 amol/ug. To demonstrate effects of both i) stromal elements and ii) clonal sub-populations within tumor masses,^14, 23, 25-28, 34^ a mixing study of *HER2*-amplified and non-amplified cell lines (OE-19/MKN-1, respectively) exemplified a dilutional effect of HER2 expression upon lowering OE-19 concentrations, likely recapitulating consequences of intra-tumoral HER2-heterogeneity, such as within CAT-14a (**Figure 2D**).

### SRM, IHC and FISH on FFPE Samples

Her2-SRM results were obtained for 139 GEC samples. Among this cohort, Her2 IHC was available for 122 samples. Among these 122 IHC cases, 51 had FISH *HER2/CEP17* ratio results, and 42 with absolute GCN scores.

### Her2-SRM Expression in FFPE Samples & Comparison to HER2 FISH

Her2-SRM levels were quantitated in 139 GEC tumors (**Supplemental Table 4**), and expression was above the LOD in 99/139 (71.2 %) samples, ranging 150-24,671amol/μg (**Figure 3A**).

**Figure 3.**
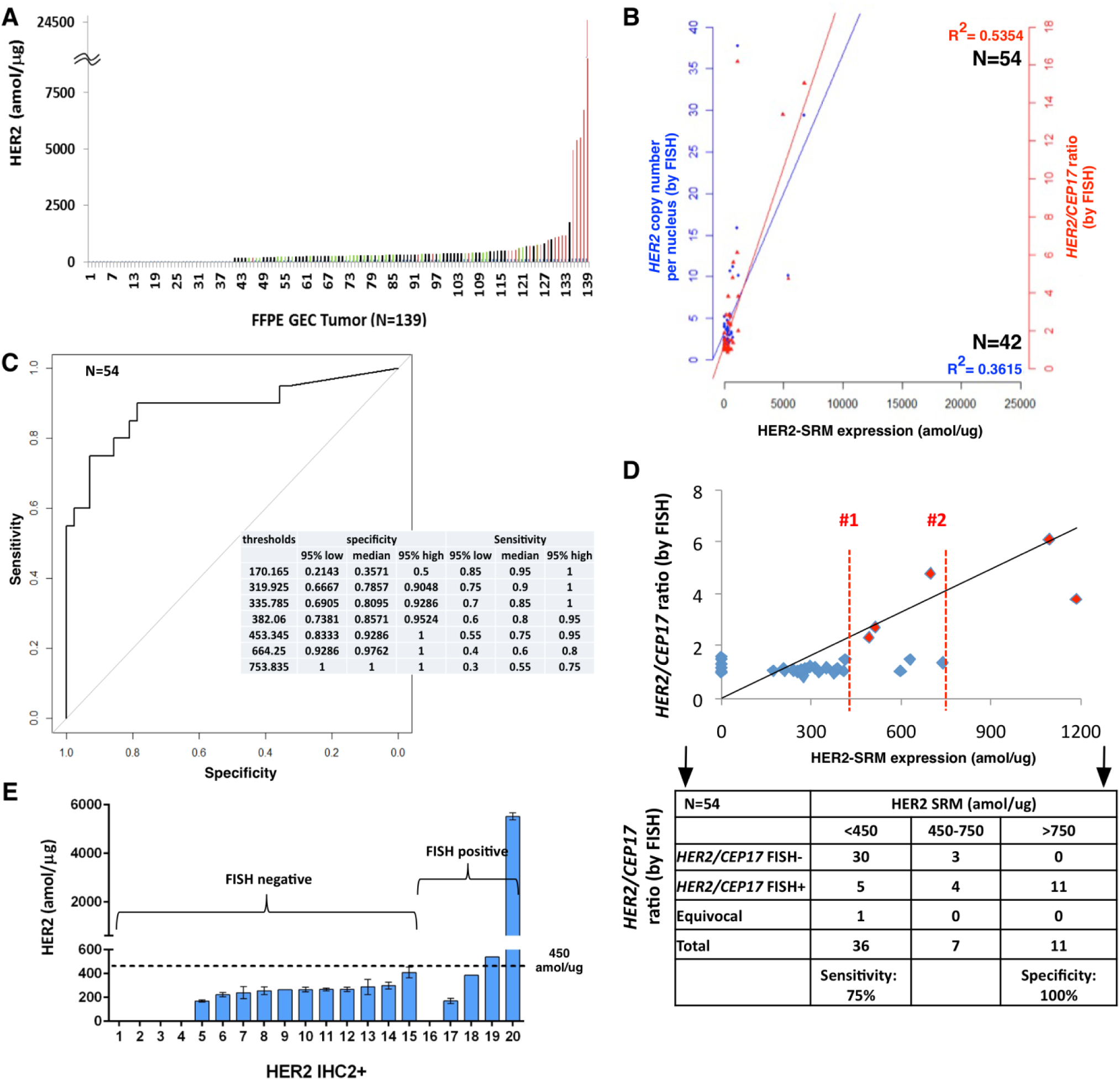
Absolute levels of Her2 in GEC tissues and correlation of Her2-SRM levels with *HER2* gene amplification. **(A)** Her2-SRM analysis of clinical FFPE GEC tissues (n=139) ranging <150-24617 amol/ug. Red highlighted samples are verified by FISH to be *HER2* amplified (*HER2/CEP17* ratio≥2); non-amplified samples are labeled as green, and samples in black were not FISH tested. **(B)** Univariate correlation of Her2-SRM and *HER2* FISH GCN (n=42, blue, r^2^ = 0.3615) and ratio (n=54, red, r^2^=0.5354). Multivariate analysis revealed stronger correlations when incorporating Met-SRM, Egfr-SRM, Her3-SRM and *HER2* FISH heterogeneity in the model: Her2-SRM:*HER2* GCN r^2^ = 0.7345 and Her2-SRM:*HER2/CEP17*ratio r^2^=0.7643 (54 Her2-SRM cases had absolute FISH ratio available). **(C)** Optimal Her2-SRM cutoff values determined by receiver operating characteristics (ROC) curve with respect to HER2/CEP17 ratio ≥2. Using one cut-off level, a value of 450 amol/ug was 75% sensitive and 93% specific to identify ‘amplification’; alternatively, a cut-off level of 750 amol/ug was 55% sensitive and 100% specific. **(D)** Using two cut-points (analogous to IHC 0/1+ = Her2 negative, and IHC 3+ = Her2 positive), with values in between (analogous to IHC 2+) = ‘equivocal’, an upper SRM level bound of 750 amol/ug and lower bound of 450 amol/ug created an equivocal range 450-750 amol/ug. The two red lines (1 and 2) represent the Her2-SRM expression falling into this equivocal range (n=9/54, 16%). (54 cases were available with binary FISH ratio data (≥2 or <2). Currently, it is recommended that these SRM-equivocal cases undergo confirmatory FISH testing. **(E)** Among tumors exhibiting Her2 IHC 2+ with FISH results (n=20), 15 tumors (75%) were FISH- and 5 (25%) were FISH+. Her2-SRM expression levels are superimposed, demonstrating that the majority (18, 90%) of these IHC2+ samples were below the 450 amol/ug SRM cut-off.

Among cases having both Her2-SRM and either FISH GCN (n=42) or FISH ratio (n=54) results, univariate analyses demonstrated fair/moderate correlation (r^2^ = 0.3615 and r^2^ = 0.5354, respectively) (**Figure 3B**). After incorporating *HER2* FISH heterogeneity along with SRM co-expression of Met, Egfr, and Her3 (Supplementary **Figure 4, Supplementary Table 5**) there was improvement in the fit of the regression model, and good correlation between HER2-SRM and FISH GCN (r^2^ = 0.7345) or FISH ratio (r^2^ = 0.7643) was observed.

**Figure 4.**
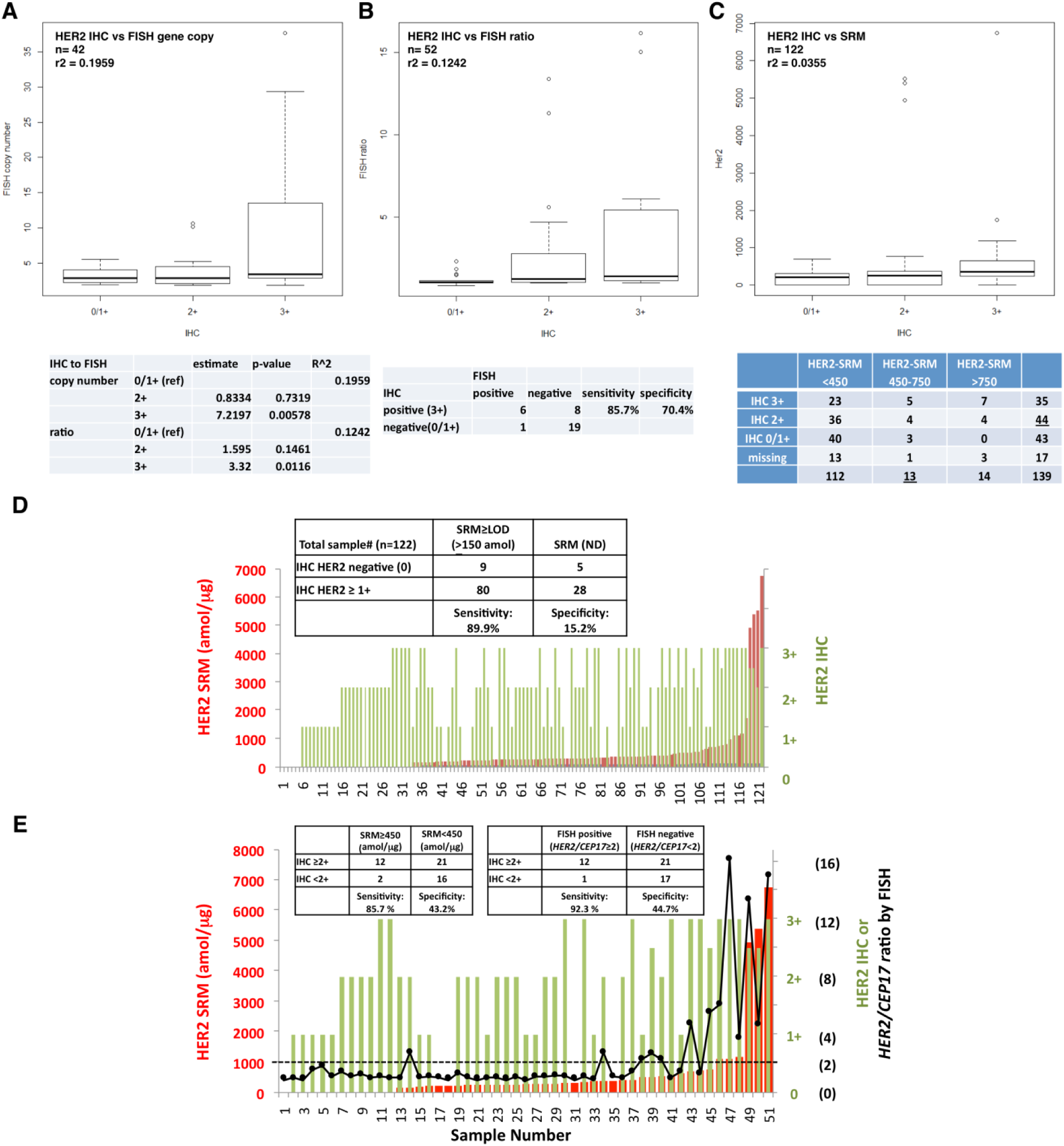
Her2 IHC correlations with FISH assay or SRM. Correlation of Her2 IHC to **(A)** FISH GCN in 42 GEC tumors (r^2^=0.1959), **(B)** *HER2/CEP17* ratio in n=52 GEC tumors (r^2^=0.1242) (52 cases with IHC results had absolute FISH ratio results); and **(C)** Her2-SRM (amol/μg) in 122 GEC tumors (r^2^=0.0355). **(D)** IHC compared to Her2-SRM with inset 2X2 table; and (E) three-way comparison of Her2-IHC, Her2-SRM and *HER2/CEP17* ratio by FISH in 51 GEC tumors where all three values were available. Inset tables demonstrate the comparison of the lower boundary for each assay (left, IHC2+ versus SRM 450amol/ug) and IHC2+ versus FISH (right). Red bar-SRM, green bar-IHC, and black dot-*HER2/CEP17* ratio by FISH. Inset tables assess sensitivity/specificity of IHC assuming SRM (D,E) and FISH (B,E) as the comparative standards.

Optimal Her2-SRM cutoffs corresponding with *HER2/CEP17* ratio ≥2 were determined using a ROC curve (**Figure 3C**). Exploring various cut-offs, 450amol/ug was 92.86% specific [95%CI 83.33-100] and 75% sensitive [95%CI 55-95] to identify *HER2* amplification by FISH; alternatively, a cut-off level of 750 amol/ug was 100% specific [95%CI 100-100] and 55% sensitive [95% CI 30-75].

While most tumors demonstrated Her2-SRM levels < 450 amol/ug (112/139 (80.1%)), a few samples had values >750 amol/μg (14/139 (10.1%)). Of 11/14 of samples >750amol/ug that were available for FISH testing, all (100%) were FISH+ (mean ratio 9.28) - a positive predictive value (PPV) of 100%. A ‘double cut-off’ level, or ‘equivocal’ zone, was applied to better identify marginal HER2 positive cases (not unlike IHC2+ ‘equivocal’). Within the identified SRM ‘equivocal zone’ of 450-750 amol/ug, there were 13 (9.4%) samples (**Figure 3C, 5B**). In terms of identifying FISH ratio≥2, the performance of the upper/lower boundaries of this equivocal zone was evaluated on the 54 cases having both Her2-SRM and FISH ratio results (**Figure 3D, 5B**). Of 7 samples (7/54, 12.9%) between 450-750 amol/ug, there were 3/7 (42.9%) that were deemed FISH- (mean FISH ratio 1.28). Therefore, the PPV was 4/7 (57.1%). A lower mean ratio (3.04) was noted for these FISH+ samples falling within the Her2-SRM equivocal zone than samples with Her2-SRM >750amol/ug (**Figure 5B**). Of 36 samples <450amol/ug (mean ratio 1.387), 5 (13.8%) were FISH+ (mean ratio 2.758), one (2.78%) was FISH ‘equivocal’, and the remaining 30 samples were FISH-, demonstrating a negative predictive value (NPV) of 83.3%. Depicting IHC, SRM and FISH results sorted by IHC explicitly demonstrates the wide range of SRM expression within the IHC 2+ and 3+ categories, and the large proportion of samples with very low SRM expression within these two groups, potentially leading to better predictive capacity with respect to benefit from anti-HER2 therapy (**Figure 5C**). In summary, using the two SRM expression boundaries, the sensitivity of the lower boundary was 75% and the specificity of the upper boundary was 100% to identify *HER2* FISH ratio≥2, and values within the Her2-SRM equivocal zone showed a FISH+ PPV of 57.1%. (**Figure 3D table**)

**Figure 5.**
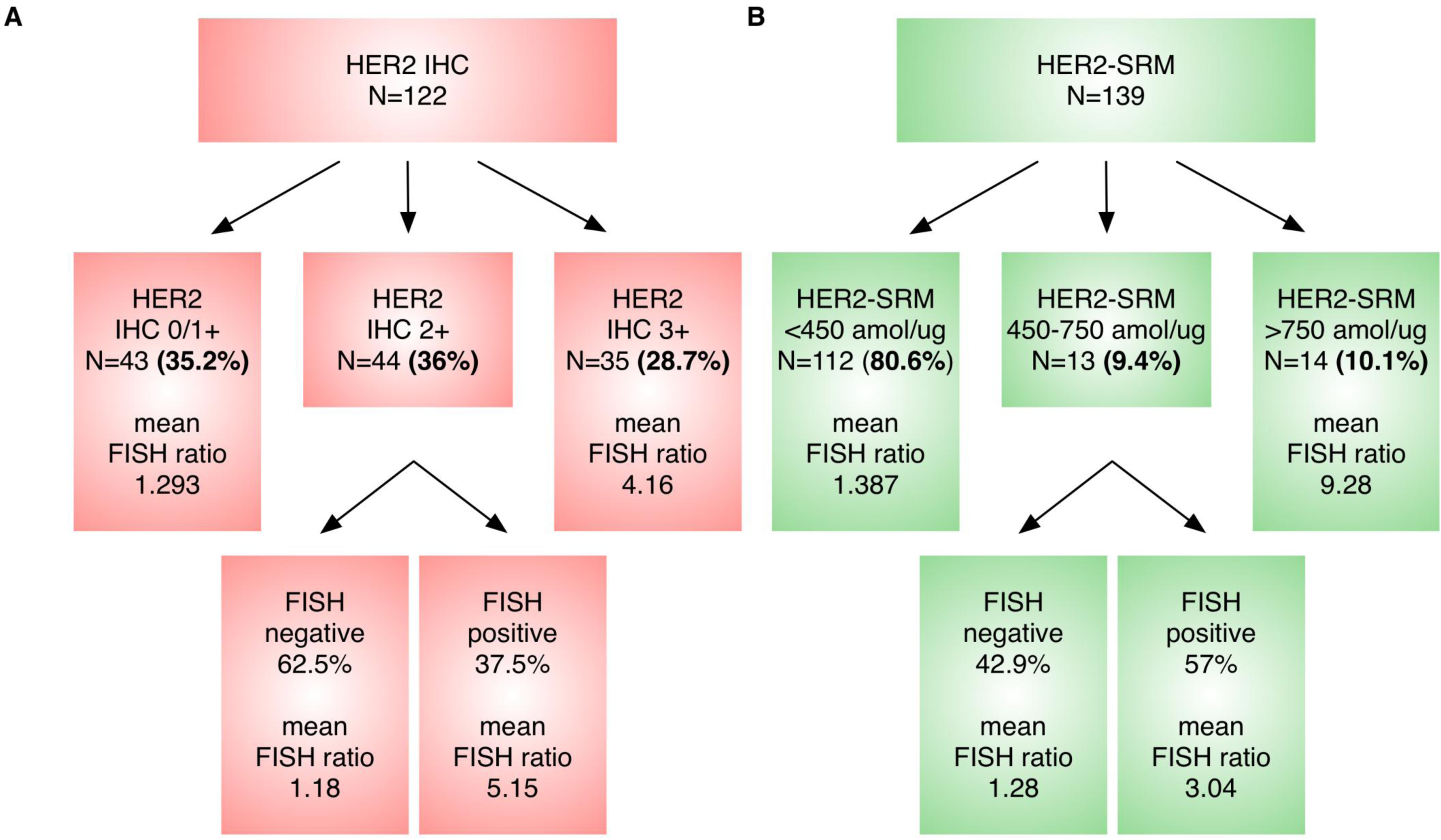
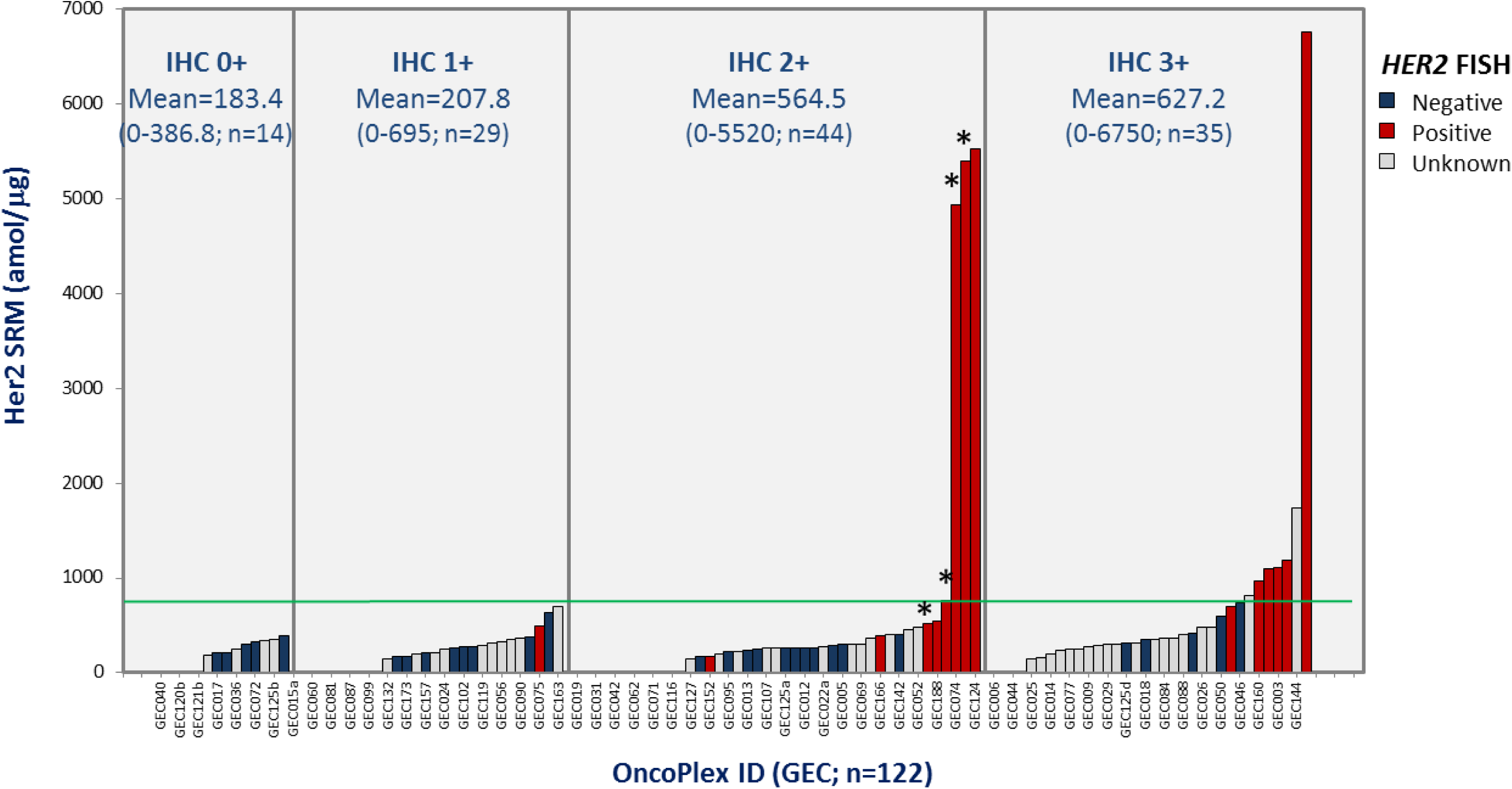
Her2 status assessment of GEC cases by IHC (A) and Her2-SRM (B) to identify HER2 ‘positive’ and ‘negative’ cases, as determined by underlying FISH *HER2/CEP17* ratio. Both assays resulted in an equivocal zone in identifying underlying *HER2* amplification (by ratio>2). However, the incidence of the Her2-SRM equivocal zone (450-750amol/ug) was much lower (9.4%) compared to IHC2+ (36%). Within these Her2-SRM equivocal cases, the PPV was relatively high (57%) compared to IHC2+ (PPV 37.5%). Her2-SRM cases >750amol/ug demonstrated a high mean *HER2/CEP17* ratio (9.28) and all were FISH amplified, compared to IHC3+ (4.16); mean ratio for Her2-SRM equivocal cases that were FISH positive was 3.04, compared to 5.15 in IHC2+/FISH+. Fewer equivocal cases requiring less reflex FISH testing along with better stratification of *HER2/CEP17* ratio demonstrated superiority of Her2-SRM over IHC. **(C)** Depicting these same IHC, SRM and FISH results now sorted by IHC category demonstrated the wide range of SRM expression within each IHC category, particularly the large proportion of samples with very low SRM expression within IHC2+ and IHC3+ groups, potentially leading to better predictive capacity of Her2-SRM with respect to benefit from anti-HER2 therapy. Four cases were scored clinically as IHC2-3+ cases (represented with ‘*) (**Supplementary Table 4**), and all four had FISH ratio >2, with three cases having Her2-SRM >750 amol/ug and one case between 450-750 amol/ug; these cases were included in the IHC 2+ category given the pathologist’s uncertainty of scoring (ie equivocal) and requirement for reflex FISH.

### Comparison of Her2 IHC2+ Status to FISH Ratio and Her2-SRM in Tissues

Among tumors having concomitant HER2 IHC/FISH/SRM testing, there were 20/54 (37%) that were IHC 2+, of which 15 (75%) were FISH- and 5 (25%) were FISH+, demonstrating a PPV of 25% (**Figure 3E**). By Her2-SRM, 18 (90%) of these IHC2+ samples were <450 amol/ug. Among all IHC samples, 44/122 (36%) were IHC2+, with PPV for FISH+ of 37.5% (**Figure 5A**). The PPV for Her2-SRM was comparatively higher (57%) within the 450-750amol/ug ‘equivocal’ zone (**Figure 3D,5B**).

### Comparison of IHC to FISH or HER2-SRM in Tissues

Among samples having both Her2 IHC and *HER2* FISH GCN (n=42) or *HER2/CEP17* FISH ratio (n=52), poor correlation was observed (r^2^=0.1959 and r^2^=0.1242, respectively) (**Figure 4A,B**). However, while IHC2+ (as referenced to IHC0/1+) was not associated with either FISH GCN or ratio, IHC3+ was significantly associated (p=0.00578 and p=0.016, respectively).

The subset of GEC samples (n=122) having both Her2 IHC (categorical variable) and Her2-SRM (linear variable) demonstrated very poor correlation, r^2^=0.0355 (**Figure 4C)**. However, when comparing IHC to SRM both as categorical variables, there was noted dependence, Chi Square p=0.02219 (**Figure 4C table**). To demonstrate the differences in sensitivity, specificity, and resolution between Her2 IHC and Her2-SRM, we compared the LODs for IHC (≥1+) versus Her2-SRM (≥150amol/ug) (**Figure 4D**). Of 122 cases, 108 (88.5%) were ≥IHC 1+, while 89 (73%) samples were ≥150amol/ug. IHC was 89.9% sensitive in identifying cases ≥150amol/ug, but only 15.2% specific in discerning SRM-negative cases. A score of ‘IHC2+’ was observed in 36% of all cases (44/122) (**Figure 5A**), or 41% (21/51) of cases available for a three-way comparison of IHC, SRM and FISH ratio. These results revealed that Her2-SRM better correlated with FISH ratio compared to IHC, with better sensitivity and specificity in identifying FISH ratio≥2 (**Figure 4E**).

### Multivariate regression model with HER2 heterogeneity and multi-plex SRM analysis of oncoproteins

To test whether the correlation between Her2-SRM expression and *HER2* FISH would improve after adjusting for the covariates ‘HER2-hetero+’, Met-SRM, Egfr-SRM, and Her3-SRM for both the cell line and tissue analyses, we evaluated a multivariate regression model (Supplementary **Figure 4A,B, Supplementary Table 5**). Interactions were observed with Her3-SRM and Egfr-SRM, (positive-interactions) as well as Met-SRM and HER2-hetero+ (negative-interactions) on the association of Her2-SRM with FISH GCN as well as with FISH ratio. These interactions were observed when either FISH status or Her2-SRM was the outcome variable.

### Clinical Correlation with Patient Vignettes

Patient 1, GEC181 (**Figure 6A**), was diagnosed clinically as stage IV HER2+ gastric cancer, yet Her2-SRM demonstrated low baseline expression (not detected, **Supplementary Table 4** sample #2) and extremely high Fgfr2-SRM. Upon rapid progression on anti-Her2 therapy, the patient responded to second line Fgfr-specific tyrosine kinase inhibition.

**Figure 6.**
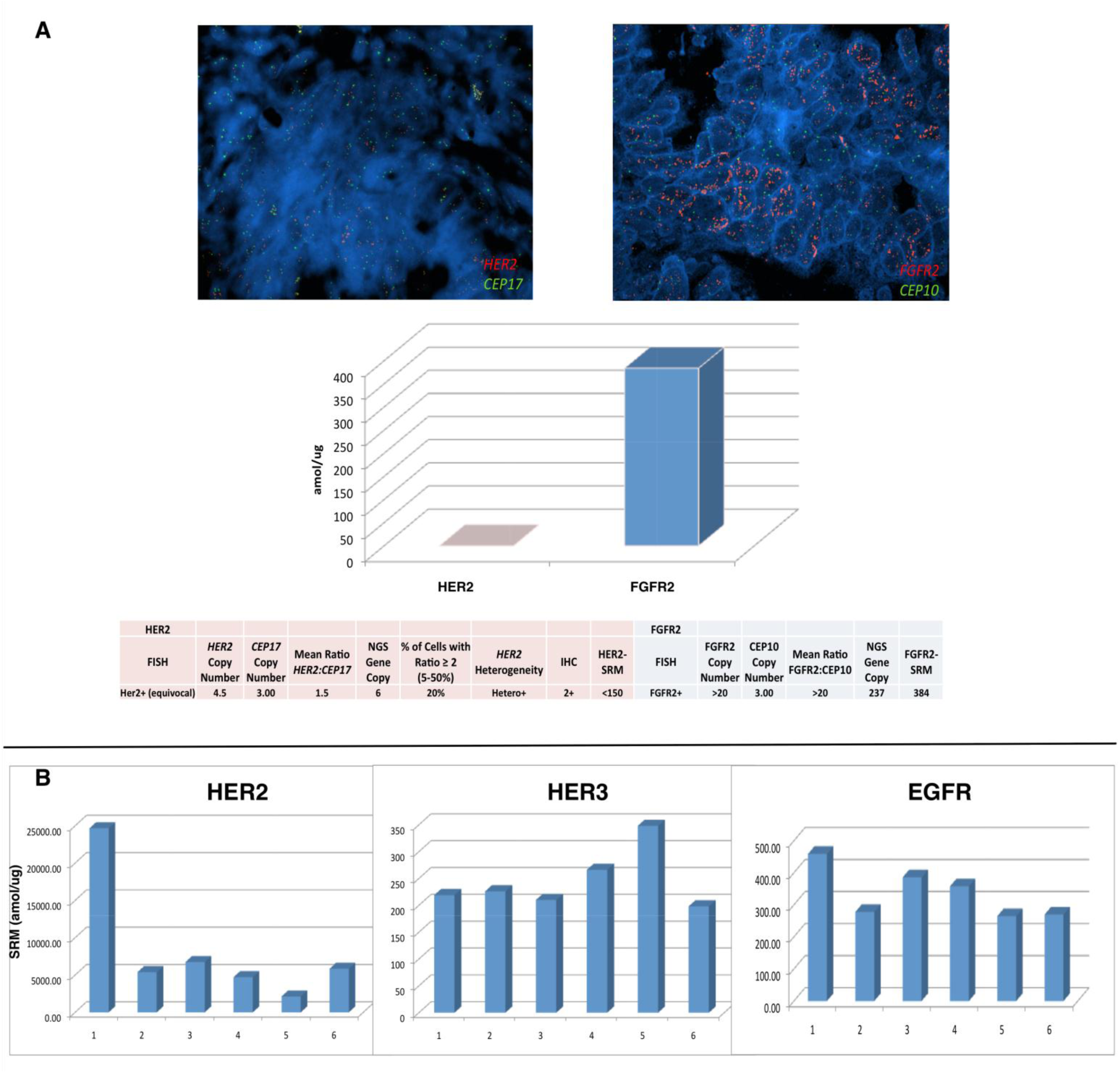
(A). Molecular profiling and a call for treatment prioritization based on *degree* of genomic/proteomic aberration. A GEC patient diagnosed clinically as ‘HER2+’ based on IHC (2+) and FISH, yet a more appropriate drug pairing was suggested by SRM multiplex testing (Fgfr2). Her2-SRM was observed to be <150 amol/ug (the range observed in all preclinical/clinical samples to date is <150- 26170 amol/ug) while Fgfr2-SRM was observed to be 384 amol/ug (above the 95 percentile of documented Fgfr2 expression). NGS, next generation sequencing. **(B) Serial Her2-SRM, Her3-SRM, and Egfr-SRM levels over time/treatment of a Her2 overexpressing and *HER2* gene amplified esophageal tumor of a 39 year old man**. **Segment 1 to 2:** First line cisplatin/irinotecan-trastuzumab 6 cycles (D1,D8 q 21 days) not well tolerated; RT to lumbar metastases; Second line FOLFIRI – trastuzumab 12 doses, SD but increasing tumor markers/bleeding; **Segment 2 to 3:** Embolization to bleeding primary tumor; Third Line –anti-PD1 antibody (PDL1+ IHC), 3 biweekly doses then PD (new multiple liver metastases); **Segment 3 to 4:** Fourth Line docetaxel-trastuzumab 10 cycles, then recurrent primary tumor bleeding but SD; **Segment 4 to 5:** RT 20Gy 1/2014-2/2014 to bleeding primary tumor. Fifth Line lapatinib/paclitaxel plus trastuzumab (15 biweekly cycles then PD on CT as well as PD clinically and by tumor markers); **Segment 5 to 6:** Sixth line FOLFOX-trastuzumab/pertuzumab – stable disease on CT with slight progression of tumor markers. **Segment 6 to present:** FOLFIRINOX-trastuzumab/pertuzumab for three cycles, response in tumors markers and PR on CT. Biopsy time points (all via EGD of primary tumor, except #6 – liver core biopsy): **1)** 5/22/2012 **2)** 6/10/2013 **3)** 10/30/2013 **4)** 1/14/2014 **5)** 10/7/2014 **6)** 12/17/15. Diagnosed with symptoms 1/2012, ultimate tissue diagnosis 5/2012 with stage IV disease (bone, M1 lymph nodes) - now almost 36 months from onset of symptoms, 33 months from biopsy (as of 2/2015).

Patient 2, GEC159 (**Figure 6B**), was diagnosed stage IV HER2+ esophago-gastric cancer, with extremely high baseline Her2-SRM levels observed (24671 amol/ug, **Supplementary Table 4** sample #139). Upon interval treatments, serial primary tumor biopsies revealed interesting SRM-expression evolutionary patterns. Upon initial trastuzumab exposure, a dramatic decrease (-78.1%) in Her2-SRM was noted at first tumor progression. After treatment with trastuzumab and lapatinib, elevation of Her3-SRM (+30.8%) was observed (+58.9% from baseline). Subsequently, addition of pertuzumab led to the most recent clinical and biochemical responses with improved dysphagia/tumor markers. The patient is maintained on this therapy 35 months from diagnosis.

## Discussion

Current HER2 diagnostics have recognized limitations, and there is urgent need for more objective, expedient, and ‘tissue economic’ assays in order to optimize clinical outcomes for patients.^8, 13, 16, 20, 23, 35^ We developed a Her2-SRM assay within a multiplex proteomic quantification assay, and demonstrated its precision, stability, and reproducibility in cell lines and clinical FFPE samples, along with correlation with IHC/FISH and other relevant oncoproteins (Her3/Egfr/Met).

We observed a wide range of Her2-SRM expression not only within the entire GEC cohort (N=139), but also *within* the subgroups of ‘IHC3+’ or ‘FISH+’ samples (ranging <150-21896.7 amol/ug). Her2-SRM correlated well with *HER2* FISH amplification status, reliably identifying highly amplified samples. The degree of *HER2* amplification (*HER2/CEP17* ratio) linearly correlated with Her2-SRM, as was previously reported,^36^ just as the absolute *HER2/CEP17* ratio was recently shown to correlate with the degree of anti-Her2 therapeutic benefit.^12^ One criticism of SRM technolgy in FFPE tissue is the low sensitivity to identify very low expression levels (ie <LOD). However, when considering gene amplified proteins, it was evident that the expression levels were dramatically higher than non-amplified expression levels, and therefore this limitation does not appear to be relevant when applying the technology to identify gene amplified tumors. We have noted this across various genes/proteins of interest.^22, 23^

Most IHC2+ cases had little Her2-SRM expression. The ‘equivocal’ zone that we defined for Her2-SRM expression (450-750 amol/ug), which lacked good correlation with *HER2/CEP17* ratio, represented approximately 10-15% of cases, substantially lower than semi-quantitative IHC2+ scoring (∼35-40%). Within respective ‘equivocal zones’, the PPV for Her2-SRM was 57% compared to 25-37.5% with IHC2+; others have demonstrated a PPV within IHC2+ as low as 13%.^36^ Our IHC2+ rate is similar to that observed in the ‘TOGA’ trial, particularly if including their IHC2+/FISH- undocumented cases. These IHC2+ rates represent the current experience in routine clinical care. Evaluating he performance of IHC versus Her2-SRM, as contrasted in **Figure 5,** demonstrated the superiority of SRM over IHC in identifying truly FISH+ positive HER2 samples, importantly relying less on reflex FISH testing.

In previous years, HER2 cut-offs for IHC and FISH rendering eligibility for anti-Her2 therapy erred towards lower thresholds, likely intending to avoid missing potential benefit of anti-Her2 therapies in patients who would otherwise be given standard cytotoxics alone. However, lack of benefit for these low-expressing subgroups is now recognized.^4, 11^ Patient 1 was clinically ‘HER2+’ (IHC2+/FISH+ equivocal), yet Her2-SRM was low (<450) - this ultimately predicted lack of benefit from anti-Her2 therapy. Although further prospective validation is required in independent datasets (which is underway), the ability to further stratify HER2 status with SRM *within* currently clinically accepted HER2+ patients will have significant treatment implications by judiciously assigning anti-HER2 therapy to those patients most likely to benefit, while sparing those likely to not benefit from both the clinical and financial toxicity of such therapy. Moreover, multiplex SRM-testing to globally survey biomarkers may allow the optimal treatments towards most-likely tumor ‘drivers’ to be administered as early as possible. In this example, very high Fgfr2 expression (consistent with *FGFR2* amplification) might have trumped borderline/low Her2 expression. Trials testing *prioritized* personalized treatment algorithms based on higher-throughput molecular profiling, including SRM-MS are ongoing.^14, 37^

Spatial intratumoral heterogeneity of FISH/IHC resulted in an observed dilutional Her2-SRM measurement, similar to the exemplary cell line mixing study. It is likely that the identified Her2-SRM ‘cut-off’ for cell lines was higher than that of tissues (>1175 versus >750 amol/ug) due to less subclonal and stromal influences. Supporting this, the CAT-14a cell line, which demonstrated HER2-heterogeneity, demonstrated Her2-SRM levels (969.33 amol/ug) lower than the 1175 amol/ug cell line cut-off. As such, the Her2-SRM assay on FFPE samples inherently captured intra-tumoral HER2 clonal heterogeneity by effectively providing an objective aggregate Her2 expression level representing all the tumor sampled via microdissection using standard H&E staining. As hypothesized, an improved correlation between Her2-SRM level and *HER2/CEP17* ratio, was observed when including the HER2-heterogeneity status into the multivariate linear regression model. The improved correlation between SRM and FISH after adjusting for FISH-heterogeneity was likely due FISH scores reflecting certain select areas of tumor (eg. invasive front, areas with higher FISH ratio than others), while SRM selects all H&E tumor indiscriminately. A significant negative interaction between Her2-SRM level and presence of *HER2*FISH heterogeneity was therefore demonstrated (lower Her2-SRM level with presence of HER2-heterogeneity).

Additionally, Her2 functional interactions with Met, Her3, and Egfr have been described. ^26, 36, 38-43^ After adjusting for these three covariates (Met/Her3/Egfr-SRM co-expression), a stronger linear correlation between Her2-SRM and FISH *HER2/CEP17* ratio was observed. Although the mechanisms for each of these interactions are not clearly defined, ultimately, gene amplification is a surrogate marker for protein overexpression. Specifically, the relationship between gene amplification and protein expression is likely multifactorial, and may be influenced by the expression of other key oncoproteins within the cell. *HER2* amplified tumors tended to have relatively lower Her2-SRM levels if they were also highly expressing SRM-Met, compared to *HER2* amplified tumors that were not highly expressing SRM-Met. Supporting our findings, Met overexpression has been linked with resistance to anti-Her2 therapy for *HER2* amplified tumors, and vice-versa.^38, 40^ On the other hand, Her3- and Egfr-SRM levels were observed to be positively associated with Her2-SRM levels, and both receptors have been implicated in signal transduction of *HER2* amplified tumors. Regardless, further work to understand these associations more clearly is required. Notwithstanding, it is possible that incorporating these co-expression covariates, such as is feasible with SRM-MS multiplex technology, while assessing clinical outcome with anti-Her2 therapies will better identify most-likely responders, and may also direct better future multi-drug targeted regimens.^38, 40, 42, 44^ This is currently being assessed prospectively in a clinically linked independent dataset.

The ability to evaluate molecular heterogeneity longitudinally through time and treatment was demonstrated using the SRM-multiplex assay in Patient 2. Extremely high Her2-SRM levels appeared to portend for prolonged benefit from anti-Her2 trastuzumab therapy (∼12 months, twice the median progression free survival in the ToGA study) before first progression (**Figure 6B** Segment 1-2). At trastuzumab-progression, Her2-SRM levels were ∼5-fold lower, offering a potential mechanism of resistance by down-regulating receptor expression - yet it remained well above IHC, FISH, and SRM cut-offs for HER2 positivity. Her2-SRM expression increased slightly after withdrawing trastuzumab for brief anti-PD1 therapy (Segment 2-3), providing more evidence of continued ‘HER2-addiction’. This expression trend, along with previous evidence that maintaining therapeutic inhibition beyond progression upon a persistent oncogenic-driver, provided rationale for resuming trastuzumab-based therapy (Segment 3-4).^14, 45-48^ After an initial response to reintroduction of trastuzumab-based therapy, but upon further progression, ‘vertical inhibition’ with lapatinib/trastuzumab led to continued response (Segment 4-5).^48, 49^ Finally, evolution towards higher Her3-SRM expression suggested another mechanism of resistance; pertuzumab-based therapy was then introduced with clinical benefit (Segments 5-6-present).^36, 43, 50^ Serial testing in order to ‘re-target’ therapies based on ‘real-time’ molecular profiles merits further testing in ongoing novel prospective clinical trial designs.^14^

Compared to IHC, SRM-MS provided more quantitative Her2 expression with better *HER2* FISH correlation, and a narrower ‘equivocal zone’. Ultimately, FISH testing for *HER2* amplification is a surrogate for Her2 protein overexpression, and we showed that this expression level is influenced by several factors, including not only *HER2/CEP17* ratio, but also HER2-heterogeneity within the sample, along with co-expression levels of various other critical oncoproteins. Therefore, a single Her2-SRM expression cut-off in the context of the SRM-MS ‘GEC-plex’ may in the future better predict anti-HER2 therapeutic benefit without any reliance on FISH or IHC. This is the subject of ongoing evaluation and validation in a large cohort of clinically-linked samples. Along with the multiplex capability of quantifying other protein biomarkers, these results demonstrate a refined Her2 expression assay for clinical application.

## Acknowledgements

DVTC would like to thank Drs. Hedy Kindler, Ravi Salgia, Funmi Olopade and Mitchell Posner for continued support.

All authors would like to thank Dr. Michael F. Press (University of Southern California) for thoughtful and critical review of this manuscript.

## References

[1] Slamon DJ, Clark GM, Wong SG, Levin WJ, Ullrich A, McGuire WL: Human breast cancer: correlation of relapse and survival with amplification of the HER-2/neu oncogene. Science 1987, 235:177–182.

[2] Natali PG, Nicotra MR, Bigotti A, Venturo I, Slamon DJ, Fendly BM, Ullrich A: Expression of the p185 encoded by HER2 oncogene in normal and transformed human tissues. Int J Cancer 1990, 45:457–161.

[3] Slamon DJ, deKernion JB, Verma IM, Cline MJ: Expression of cellular oncogenes in human malignancies. Science 1984, 224:256–162.

[4] Bang YJ, Van Cutsem E, Feyereislova A, Chung HC, Shen L, Sawaki A, Lordick F, Ohtsu A, Omuro Y, Satoh T, Aprile G, Kulikov E, Hill J, Lehle M, Ruschoff J, Kang YK: Trastuzumab in combination with chemotherapy versus chemotherapy alone for treatment of HER2-positive advanced gastric or gastrooesophageal junction cancer (ToGA): a phase 3, open-label, randomised controlled trial. Lancet 2010, 376:687–197.

[5] Sehdev A, Catenacci DV: Gastroesophageal cancer: focus on epidemiology, classification, and staging. Discov Med 2013, 16:103–111.

[6] Hofmann M, Stoss O, Shi D, Buttner R, van de Vijver M, Kim W, Ochiai A, Ruschoff J, Henkel T: Assessment of a HER2 scoring system for gastric cancer: results from a validation study. Histopathology 2008, 52:797–1805.

[7] Ruschoff J, Dietel M, Baretton G, Arbogast S, Walch A, Monges G, Chenard MP, Penault-Llorca F, Nagelmeier I, Schlake W, Hofler H, Kreipe HH: HER2 diagnostics in gastric cancer-guideline validation and development of standardized immunohistochemical testing. Virchows Arch 2010, 457:299–1307.

[8] Ruschoff J, Hanna W, Bilous M, Hofmann M, Osamura RY, Penault-Llorca F, van de Vijver M, Viale G: HER2 testing in gastric cancer: a practical approach. Mod Pathol 2012, 25:637–150.

[9] Bartley AN, Christ J, Fitzgibbons PL, Hamilton SR, Kakar S, Shah MA, Tang LH, Troxell ML: Template for Reporting Results of HER2 (ERBB2) Biomarker Testing of Specimens From Patients With Adenocarcinoma of the Stomach or Esophagogastric Junction. Arch Pathol Lab Med 2014.

[10] Hecht JR, Bang YJ, Qin S, Chung HC, Xu JM, Park JO, Jeziorski K: Lapatinib in combination with capecitabine plus oxaliplatin (CapeOx) in HER2-positive advanced or metastatic gastric, esophgael, or gastroesophageal adenocarcinoma (AC): The TRIO-013/LOGiC Trial. J Clin Oncol 2013, 31, abstr LBA4001.

[11] Satoh T, Xu RH, Chung HC, Sun GP, Doi T, Xu JM, Tsuji A, Omuro Y, Li J, Wang JW, Miwa H, Qin SK, Chung IJ, Yeh KH, Feng JF, Mukaiyama A, Kobayashi M, Ohtsu A, Bang YJ: Lapatinib plus paclitaxel versus paclitaxel alone in the second-line treatment of HER2-amplified advanced gastric cancer in Asian populations: TyTAN-a randomized, phase III study. J Clin Oncol 2014, 32:2039–149.

[12] Gomez-Martin C, Plaza JC, Pazo-Cid R, Salud A, Pons F, Fonseca P, Leon A, Alsina M, Visa L, Rivera F, Galan MC, Del Valle E, Vilardell F, Iglesias M, Fernandez S, Landolfi S, Cuatrecasas M, Mayorga M, Jose Paules M, Sanz-Moncasi P, Montagut C, Garralda E, Rojo F, Hidalgo M, Lopez-Rios F: Level of HER2 gene amplification predicts response and overall survival in HER2-positive advanced gastric cancer treated with trastuzumab. J Clin Oncol 2013, 31:4445–152.

[13] Khoury JD, Catenacci DV: Next-Generation Companion Diagnostics: Promises, Challenges, and Solutions. Arch Pathol Lab Med 2014.

[14] Catenacci DVT: Next-generation clinical trials: Novel strategies to address the challenge of tumor molecular heterogeneity. Molecular Oncology 2014.

[15] Allison M: The HER2 testing conundrum. Nat Biotechnol 2010, 28:117–19.

[16] Carlson B: HER2 TESTS: How Do We Choose? Biotechnol Healthc 2008, 5:23–17.

[17] Cho EY, Srivastava A, Park K, Kim J, Lee MH, Do I, Lee J, Kim KM, Sohn TS, Kang WK, Kim S: Comparison of four immunohistochemical tests and FISH for measuring HER2 expression in gastric carcinomas. Pathology 2012, 44:216–120.

[18] Buza N, English DP, Santin AD, Hui P: Toward standard HER2 testing of endometrial serous carcinoma: 4-year experience at a large academic center and recommendations for clinical practice. Mod Pathol 2013, 26:1605–112.

[19] McCullough AE, Dell’orto P, Reinholz MM, Gelber RD, Dueck AC, Russo L, Jenkins RB, Andrighetto S, Chen B, Jackisch C, Untch M, Perez EA, Piccart-Gebhart MJ, Viale G: Central pathology laboratory review of HER2 and ER in early breast cancer: an ALTTO trial [BIG 2-06/NCCTG N063D (Alliance)] ring study. Breast Cancer Res Treat 2014, 143:485–192.

[20] O’Hurley G, Sjostedt E, Rahman A, Li B, Kampf C, Ponten F, Gallagher WM, Lindskog C: Garbage in, garbage out: a critical evaluation of strategies used for validation of immunohistochemical biomarkers. Mol Oncol 2014, 8:783–198.

[21] Hembrough T, Thyparambil S, Liao WL, Darfler MM, Abdo J, Bengali KM, Hewitt SM, Bender RA, Krizman DB, Burrows J: Application of selected reaction monitoring for multiplex quantification of clinically validated biomarkers in formalin-fixed, paraffin-embedded tumor tissue. J Mol Diagn 2013, 15:454–165.

[22] Hembrough T, Thyparambil S, Liao WL, Darfler MM, Abdo J, Bengali KM, Taylor P, Tong J, Lara-Guerra H, Waddell TK, Moran MF, Tsao MS, Krizman DB, Burrows J: Selected Reaction Monitoring (SRM) Analysis of Epidermal Growth Factor Receptor (EGFR) in Formalin Fixed Tumor Tissue. Clin Proteomics 2012, 9:5.

[23] Catenacci DV, Liao WL, Thyparambil S, Henderson L, Xu P, Zhao L, Rambo B, Hart J, Xiao SY, Bengali K, Uzzell J, Darfler M, Krizman DB, Cecchi F, Bottaro DP, Karrison T, Veenstra TD, Hembrough T, Burrows J: Absolute quantitation of Met using mass spectrometry for clinical application: assay precision, stability, and correlation with MET gene amplification in FFPE tumor tissue. PLoS One 2014, 9:e100586.

[24] Hembrough T, Henderson L, Rambo B, Liao WL, Thyparambil S, Bengali K, Uzzell J, Darfler M, Krizman DB, Xu P, Xiao SY, Zhao L, Burrows J, Catenacci DV: Quantification of HER2 from Gastroesophageal Cancer (GEC) FFPE Tissue by Mass Spectrometry (MS). J Clin Oncol 32, 2014 (suppl 3; abstr 17).

[25] Seol H, Lee HJ, Choi Y, Lee HE, Kim YJ, Kim JH, Kang E, Kim SW, Park SY: Intratumoral heterogeneity of HER2 gene amplification in breast cancer: its clinicopathological significance. Mod Pathol 2012, 25:938–148.

[26] Lee HE, Park KU, Yoo SB, Nam SK, Park do J, Kim HH, Lee HS: Clinical significance of intratumoral HER2 heterogeneity in gastric cancer. Eur J Cancer 2013, 49:1448–157.

[27] Arena V, Pennacchia I, Vecchio FM, Carbone A: HER-2 intratumoral heterogeneity. Mod Pathol 2013, 26:607–19.

[28] Lee HJ, Park SY: Reply to ‘Intratumoral heterogeneity of HER2 gene amplification in breast cancer: its clinicopathological significance’. Mod Pathol 2013, 26:610–11.

[29] Nilsson T, Mann M, Aebersold R, Yates JR, 3rd, Bairoch A, Bergeron JJ: Mass spectrometry in high-throughput proteomics: ready for the big time. Nat Methods 2010, 7:681–15.

[30] Addona TA, Abbatiello SE, Schilling B, Skates SJ, Mani DR, Bunk DM, Spiegelman CH, Zimmerman LJ, Ham AJ, Keshishian H, Hall SC, Allen S, Blackman RK, Borchers CH, Buck C, Cardasis HL, Cusack MP, Dodder NG, Gibson BW, Held JM, Hiltke T, Jackson A, Johansen EB, Kinsinger CR, Li J, Mesri M, Neubert TA, Niles RK, Pulsipher TC, Ransohoff D, Rodriguez H, Rudnick PA, Smith D, Tabb DL, Tegeler TJ, Variyath AM, Vega-Montoto LJ, Wahlander A, Waldemarson S, Wang M, Whiteaker JR, Zhao L, Anderson NL, Fisher SJ, Liebler DC, Paulovich AG, Regnier FE, Tempst P, Carr SA: Multi-site assessment of the precision and reproducibility of multiple reaction monitoring-based measurements of proteins in plasma. Nat Biotechnol 2009, 27:633–41.

[31] Whiteaker JR, Lin C, Kennedy J, Hou L, Trute M, Sokal I, Yan P, Schoenherr RM, Zhao L, Voytovich UJ, Kelly-Spratt KS, Krasnoselsky A, Gafken PR, Hogan JM, Jones LA, Wang P, Amon L, Chodosh LA, Nelson PS, McIntosh MW, Kemp CJ, Paulovich AG: A targeted proteomics-based pipeline for verification of biomarkers in plasma. Nat Biotechnol 2011, 29:625–134.

[32] Catenacci DV, Cervantes G, Yala S, Nelson EA, El-Hashani E, Kanteti R, El Dinali M, Hasina R, Bragelmann J, Seiwert T, Sanicola M, Henderson L, Grushko TA, Olopade O, Karrison T, Bang YJ, Kim WH, Tretiakova M, Vokes E, Frank DA, Kindler HL, Huet H, Salgia R: RON (MST1R) is a novel prognostic marker and therapeutic target for gastroesophageal adenocarcinoma. Cancer Biol Ther 2011, 12:9–146.

[33] Catenacci DV, Henderson L, Xiao SY, Patel P, Yauch RL, Hegde P, Zha J, Pandita A, Peterson A, Salgia R: Durable complete response of metastatic gastric cancer with anti-Met therapy followed by resistance at recurrence. Cancer Discov 2011, 1:573–19.

[34] Catenacci D, Polite B, Henderson L, Xu P, Rambo B, Liao WL, Hembrough T, Burrows J, Zhao L, Hart J, Xiao SY, Karrison T, Dignam J, Kinder HL: Towards personalized treatment for gastroesophageal adenocarcinoma (GEC): Strategies to address tumor heterogeneity - PANGEA. J Clin Oncol 2014, GI ASCO 2014, Abstr 60.

[35] Chia S: Testing for discordance at metastatic relapse: does it matter? J Clin Oncol 2012, 30:575–16.

[36] Yoon HH, Sukov WR, Shi Q, Sattler CA, Wiktor AE, Diasio RB, Wu TT, Jenkins RB, Sinicrope FA: HER-2/neu gene amplification in relation to expression of HER2 and HER3 proteins in patients with esophageal adenocarcinoma. Cancer 2014, 120:415–124.

[37] Abrams J, Conley B, Mooney M, Zwiebel J, Chen A, Welch JJ, Takebe N, Malik S, McShane L, Korn E, Williams M, Staudt L, Doroshow J: National Cancer Institute’s Precision Medicine Initiatives for the new National Clinical Trials Network. Am Soc Clin Oncol Educ Book 2014:71–16.

[38] Chen CT, Kim H, Liska D, Gao S, Christensen JG, Weiser MR: MET activation mediates resistance to lapatinib inhibition of HER2-amplified gastric cancer cells. Mol Cancer Ther 2012, 11:660–19.

[39] Yonesaka K, Zejnullahu K, Okamoto I, Satoh T, Cappuzzo F, Souglakos J, Ercan D, Rogers A, Roncalli M, Takeda M, Fujisaka Y, Philips J, Shimizu T, Maenishi O, Cho Y, Sun J, Destro A, Taira K, Takeda K, Okabe T, Swanson J, Itoh H, Takada M, Lifshits E, Okuno K, Engelman JA, Shivdasani RA, Nishio K, Fukuoka M, Varella-Garcia M, Nakagawa K, Janne PA: Activation of ERBB2 signaling causes resistance to the EGFR-directed therapeutic antibody cetuximab. Sci Transl Med 2011, 3:99ra86.

[40] Paulson AK, Linklater ES, Berghuis BD, App CA, Oostendorp LD, Paulson JE, Pettinga JE, Melnik MK, Vande Woude GF, Graveel CR: MET and ERBB2 are coexpressed in ERBB2+ breast cancer and contribute to innate resistance. Mol Cancer Res 2013, 11:1112–121.

[41] Shattuck DL, Miller JK, Laederich M, Funes M, Petersen H, Carraway KL, 3rd, Sweeney C: LRIG1 is a novel negative regulator of the Met receptor and opposes Met and Her2 synergy. Mol Cell Biol 2007, 27:1934–146.

[42] Rusnak DW, Alligood KJ, Mullin RJ, Spehar GM, Arenas-Elliott C, Martin AM, Degenhardt Y, Rudolph SK, Haws TF, Jr., Hudson-Curtis BL, Gilmer TM: Assessment of epidermal growth factor receptor (EGFR, ErbB1) and HER2 (ErbB2) protein expression levels and response to lapatinib (Tykerb, GW572016) in an expanded panel of human normal and tumour cell lines. Cell Prolif 2007, 40:580–194.

[43] Zhang XL, Yang YS, Xu DP, Qu JH, Guo MZ, Gong Y, Huang J: Comparative study on overexpression of HER2/neu and HER3 in gastric cancer. World J Surg 2009, 33:2112–18.

[44] Yoon HH, Shi Q, Sukov WR, Lewis MA, Sattler CA, Wiktor AE, Wu TT, Diasio RB, Jenkins RB, Sinicrope FA: Adverse prognostic impact of intratumor heterogeneous HER2 gene amplification in patients with esophageal adenocarcinoma. Journal of clinical oncology: official journal of the American Society of Clinical Oncology 2012, 30:3932–18.

[45] Verma S, Miles D, Gianni L, Krop IE, Welslau M, Baselga J, Pegram M, Oh DY, Dieras V, Guardino E, Fang L, Lu MW, Olsen S, Blackwell K: Trastuzumab emtansine for HER2-positive advanced breast cancer. N Engl J Med 2012, 367:1783–191.

[46] von Minckwitz G, du Bois A, Schmidt M, Maass N, Cufer T, de Jongh FE, Maartense E, Zielinski C, Kaufmann M, Bauer W, Baumann KH, Clemens MR, Duerr R, Uleer C, Andersson M, Stein RC, Nekljudova V, Loibl S: Trastuzumab beyond progression in human epidermal growth factor receptor 2-positive advanced breast cancer: a german breast group 26/breast international group 03-05 study. J Clin Oncol 2009, 27:1999–12006.

[47] Geyer CE, Forster J, Lindquist D, Chan S, Romieu CG, Pienkowski T, Jagiello-Gruszfeld A, Crown J, Chan A, Kaufman B, Skarlos D, Campone M, Davidson N, Berger M, Oliva C, Rubin SD, Stein S, Cameron D: Lapatinib plus capecitabine for HER2-positive advanced breast cancer. N Engl J Med 2006, 355:2733–143.

[48] Blackwell KL, Burstein HJ, Storniolo AM, Rugo HS, Sledge G, Aktan G, Ellis C, Florance A, Vukelja S, Bischoff J, Baselga J, O’Shaughnessy J: Overall survival benefit with lapatinib in combination with trastuzumab for patients with human epidermal growth factor receptor 2-positive metastatic breast cancer: final results from the EGF104900 Study. J Clin Oncol 2012, 30:2585–192.

[49] Moasser MM: Two dimensions in targeting HER2. J Clin Oncol 2014, 32:2074–17.

[50] Baselga J, Cortes J, Kim SB, Im SA, Hegg R, Im YH, Roman L, Pedrini JL, Pienkowski T, Knott A, Clark E, Benyunes MC, Ross G, Swain SM: Pertuzumab plus trastuzumab plus docetaxel for metastatic breast cancer. N Engl J Med 2012, 366:109–119.

